# Effects of HMGCR deficiency on skeletal muscle development

**DOI:** 10.1101/2024.05.06.591934

**Authors:** Mekala Gunasekaran, Hannah R. Littel, Natalya M. Wells, Johnnie Turner, Gloriana Campos, Sree Venigalla, Elicia A. Estrella, Partha S. Ghosh, Audrey L. Daugherty, Seth A. Stafki, Louis M. Kunkel, A. Reghan Foley, Sandra Donkervoort, Carsten G. Bönnemann, Laura Toledo-Bravo de Laguna, Andres Nascimento, Daniel Natera-de Benito, Isabelle Draper, Christine C. Bruels, Christina A. Pacak, Peter B. Kang

**Affiliations:** Greg Marzolf Jr. Muscular Dystrophy Center and Department of Neurology, University of Minnesota Medical School, Minneapolis, Minnesota, USA; Division of Genetics and Genomics, Boston Children’s Hospital and Harvard Medical School, Boston, Massachusetts, USA; Department of Neurology, Boston Children’s Hospital and Harvard Medical School, Boston, Massachusetts, USA; Neuromuscular and Neurogenetic Disorders of Childhood Section, Neurogenetics Branch, National Institute of Neurological Disorders and Stroke, NIH, Bethesda, MD, USA; Department of Pediatrics, Hospital Materno-Infantil, Las Palmas de Gran Canaria, Spain; Neuromuscular Unit, Department of Neurology, Hospital Sant Joan de Déu, Barcelona, Spain; Applied Research in Neuromuscular Diseases, Institut de Recerca Sant Joan de Déu, Barcelona, Spain; Molecular Cardiology Research Institute, Tufts Medical Center, Boston, Massachusetts, USA; Institute for Translational Neuroscience, University of Minnesota, Minneapolis, Minnesota, USA

**Keywords:** HMGCR, muscular dystrophy, myoblast, skeletal muscle

## Abstract

Pathogenic variants in *HMGCR* were recently linked to a limb-girdle muscular dystrophy (LGMD) phenotype. The protein product HMG CoA reductase (HMGCR) catalyzes a key component of the cholesterol synthesis pathway. The two other muscle diseases associated with HMGCR, statin-associated myopathy (SAM) and autoimmune anti-HMGCR myopathy, are not inherited in a Mendelian pattern. The mechanism linking pathogenic variants in *HMGCR* with skeletal muscle dysfunction is unclear. We knocked down *Hmgcr* in mouse skeletal myoblasts, knocked down *hmgcr* in Drosophila, and expressed three pathogenic *HMGCR* variants (c.1327C>T, p.Arg443Trp; c.1522_1524delTCT, p.Ser508del; and c.1621G>A, p.Ala541Thr) in *Hmgcr* knockdown mouse myoblasts. *Hmgcr* deficiency was associated with decreased proliferation, increased apoptosis, and impaired myotube fusion. Transcriptome sequencing of *Hmgcr* knockdown versus control myoblasts revealed differential expression involving mitochondrial function, with corresponding differences in cellular oxygen consumption rates. Both ubiquitous and muscle-specific knockdown of *hmgcr* in Drosophila led to lethality. Overexpression of reference *HMGCR* cDNA rescued myotube fusion in knockdown cells, whereas overexpression of the pathogenic variants of *HMGCR* cDNA did not. These results suggest that the three HMGCR-related muscle diseases share disease mechanisms related to skeletal muscle development.

## Introduction

HMGCR is a vital rate-limiting glycoprotein enzyme within the mevalonate and isoprenoid pathways that converts 3-hydroxy-3-methy-glutaryl coenzyme A (HMG-CoA) to mevalonate (Figure 1). This reaction is an essential step leading to the cholesterol synthesis pathway, including squalene mediated cholesterol synthesis and geranylgeranyl pyrophosphate mediated protein prenylation^1,2^. Apart from these, mevalonate is also a precursor of dolichol, which mediates glycoprotein synthesis, and the anti-oxidant coenzyme Q10 (also called ubiquinone) which is essential for regulating mitochondrial function in cellular growth and maintenance^3^. Liver specific *Hmgcr* knockout is fatal in mice and this phenotype is rescued with mevalonate administration^4^. Hence HMGCR and the mevalonate pathway are therapeutic targets for certain conditions including hyperlipidemia, hypercholesterolemia, and cancer^5–8^. The most widely used HMGCR targeting medications are statins, which inhibit HMGCR activity. Statin-associated myopathy (SAM) is a common and sometimes debilitating side effect of statin therapy^9,10^. Additionally, a small proportion of individuals taking statins develop an immune-mediated necrotizing muscle disease, known as anti-HMGCR myopathy^11,12^, linking HMGCR inhibition to a muscle disease^13,14^ that has clinical overlap with limb-girdle muscular dystrophy (LGMD)^15^.

**Figure 1.**
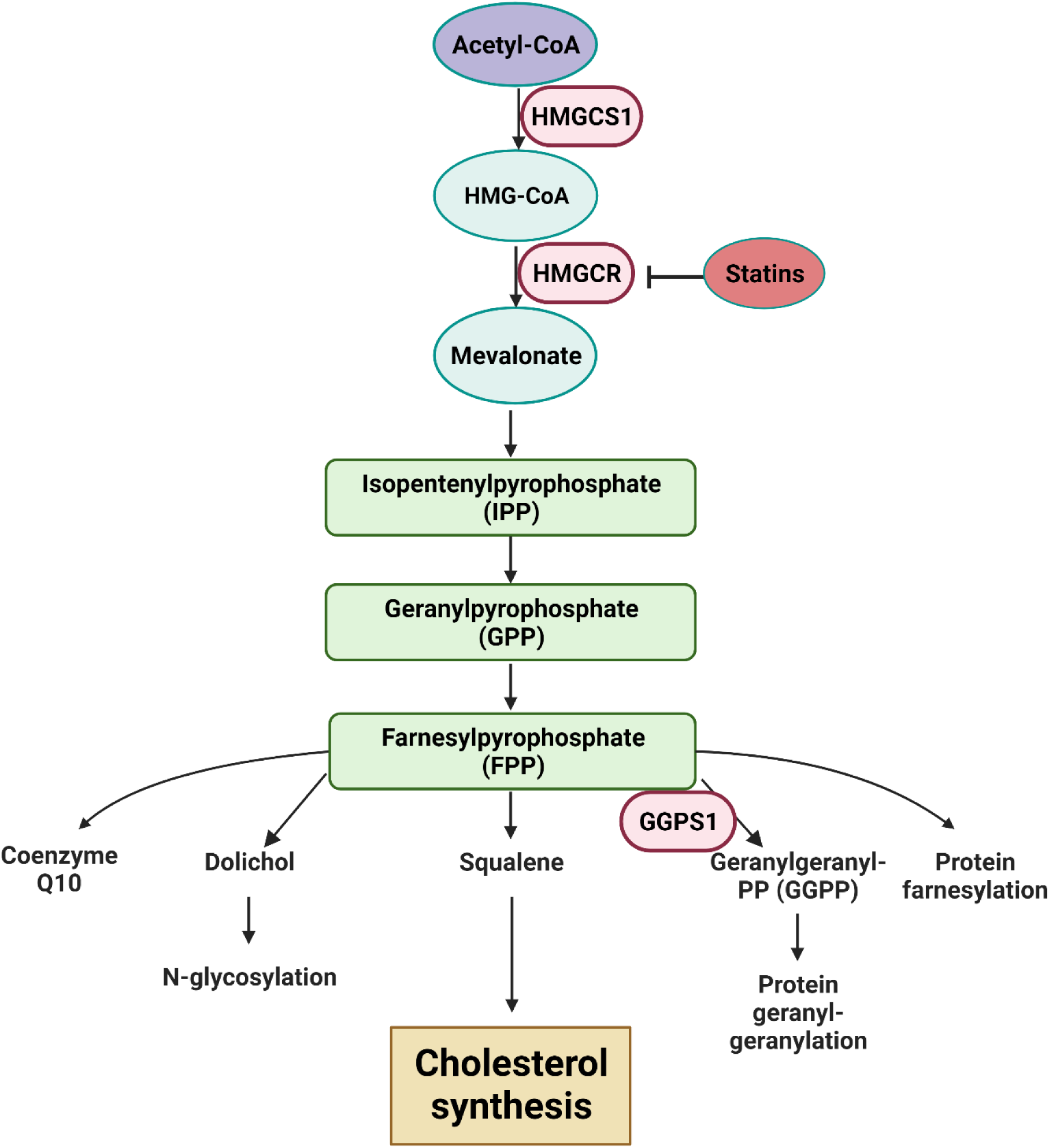
Diagram of the HMGCR metabolic pathway. 3-hydroxy-3-methylglutaryl-CoA (HMG-CoA) is converted to mevalonate by HMG-CoA reductase (HMGCR). Mevalonate then serves as a substrate for several downstream reactions, including cholesterol synthesis. Statins inhibitHMGCR, lowering cholesterol synthesis in hypercholesterolemia. Two additional enzymes associated with inherited skeletal muscle or musculoskeletal diseases are shown: HMGCS1 and GGPS1. Diagram was generated using BioRender under a license permitting its use in publications.

Two recent reports characterize a total of 6 families with biallelic pathogenic variants in *HMGCR* in the setting of LGMD phenotypes, with one individual deriving therapeutic benefit from mevalonolactone administration^16,17^. However, details of the connection between pathogenic variants in *HMGCR* and inherited muscle disease remain unclear. HSA-Cre-mediated skeletal muscle-specific knockout of *Hmgcr* in mice leads to decreased locomotor activity and muscle fibers exhibiting rhabdomyolysis with mitochondrial dysfunction^18^. We examined the effects of HMGCR deficiency in myoblast cultures and Drosophila, along with the potential rescue effects of reference and pathogenic *HMGCR* variants in myoblast cultures, providing additional functional context for HMGCR-related muscular dystrophy.

## Results

### Affected families and selection of human pathogenic variants for modeling

We ascertained three families with HMGCR-related muscular dystrophy (Figure 2a). The first family was recently published and included two siblings with severe muscle phenotypes who harbored compound heterozygous pathogenic variants in *HMGCR* (NM_000859.3: c.1327C>T, p.Arg443Trp and NM_000859.3: c.1522_1524delTCT, p.Ser508del)^16^. These variants localize to the linker and cytoplasmic catalytic domains of the protein product. A girl in the second family presented with delayed walking (28 months), exercise intolerance, an inability to run, and dysphagia.

**Figure 2.**
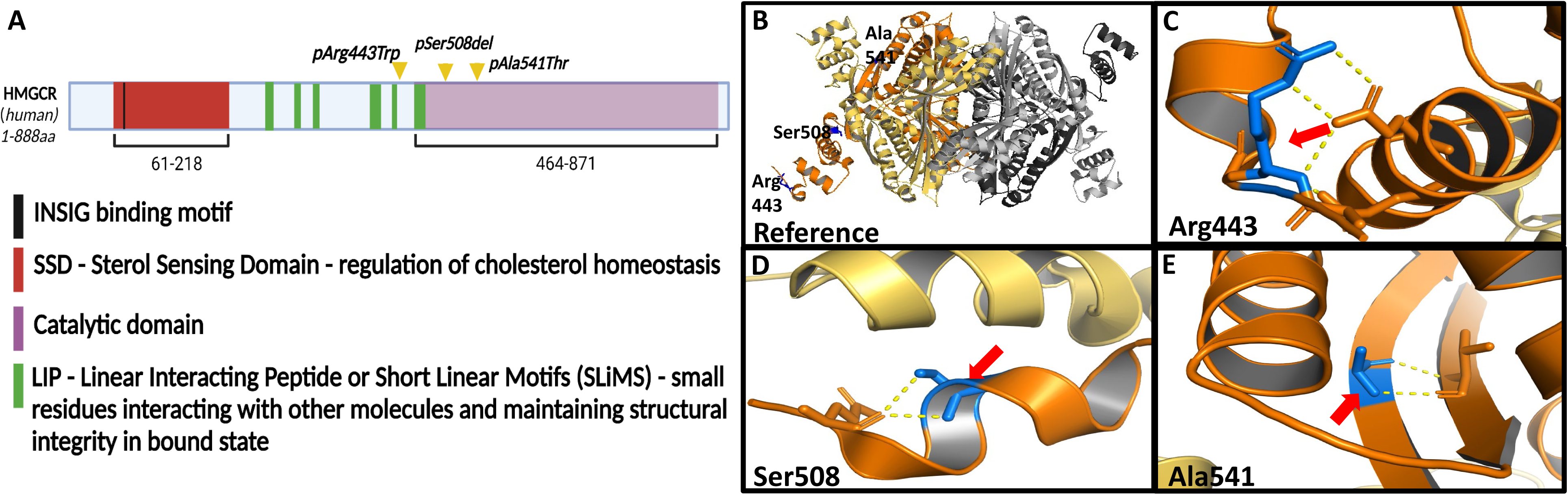
(A) Diagram of the protein domains of human HMGCR. Positions of the 3 pathogenic variants in the current study are marked with yellow arrow heads. The p.Arg443 residue is located outside the catalytic domain whereas p.Ser508 and p.Ala541 are located in the catalytic domain. Diagram was generated using BioRender under a license permitting their use in publications. (B) Diagram of the full HMGCR protein is shown with the three variants of interest. Insets of HMGCR are shown with the (C) p.Arg443, (D) p.Ser508, and € p.Ala541 residues noted. The amino acid residues of interest are colored blue and marked by red arrows. Diagrams (B-E) were generated using Pymol 2.5.0.

Neurocognitive development was unremarkable. She was found to have a homozygous pathogenic variant in *HMGCR* (NM_000859.3: c.1621G>A, p.Ala541Thr), also located in the catalytic domain of the HMGCR protein. A boy in the third family presented at the age of 3 years with elevated serum transaminase levels (1,711 U/L) which were incidentally discovered during a blood test conducted as part of the investigation for recurrent episodes of emesis. While his psychomotor development had been normal and he began walking at the age of 15 months, at age 3 years he was found to have difficulty rising from the floor and difficulty ascending stairs. Subsequent genetic analysis revealed that he had a homozygous variant in *HMGCR* (NM_000859.3: c.1522_1524delTCT, p.Ser508del). At age 5 years he was found to have neck flexor and axial weakness, and proximal limb weakness without calf pseudohypertrophy. Neurocognitive development was unremarkable. Diagrams of the positions of the pathogenic variants for all three families are shown in Figure 2b.

### shRNA mediated knockdown of Hmgcr

shRNA mediated knockdown of *Hmgcr* was performed on C2C12 myoblasts. On qPCR, shRNA knockdown myoblasts showed a ∼60% reduction in *Hmgcr* levels versus the levels in scrambled control myoblasts, confirming knockdown efficiency (Figure S1a). GFP+ clones containing shRNA-transfected cells were expanded in culture to generate stable shRNA positive cell lines (Figure S1b).

### Myoblast proliferation and apoptosis patterns in Hmgcr deficiency

*Hmgcr* knockdown impaired cell proliferation, with a significant loss of cell abundance on days 1, 2 and 3 of proliferation for *Hmgcr* knockdown cells compared to scrambled shRNA controls (Figure 3a). TUNEL assays showed 13.2% apoptotic cells in the setting of *Hmgcr* knockdown compared to 5.7% in scrambled controls (Figure 3b & 3c).

**Figure 3.**
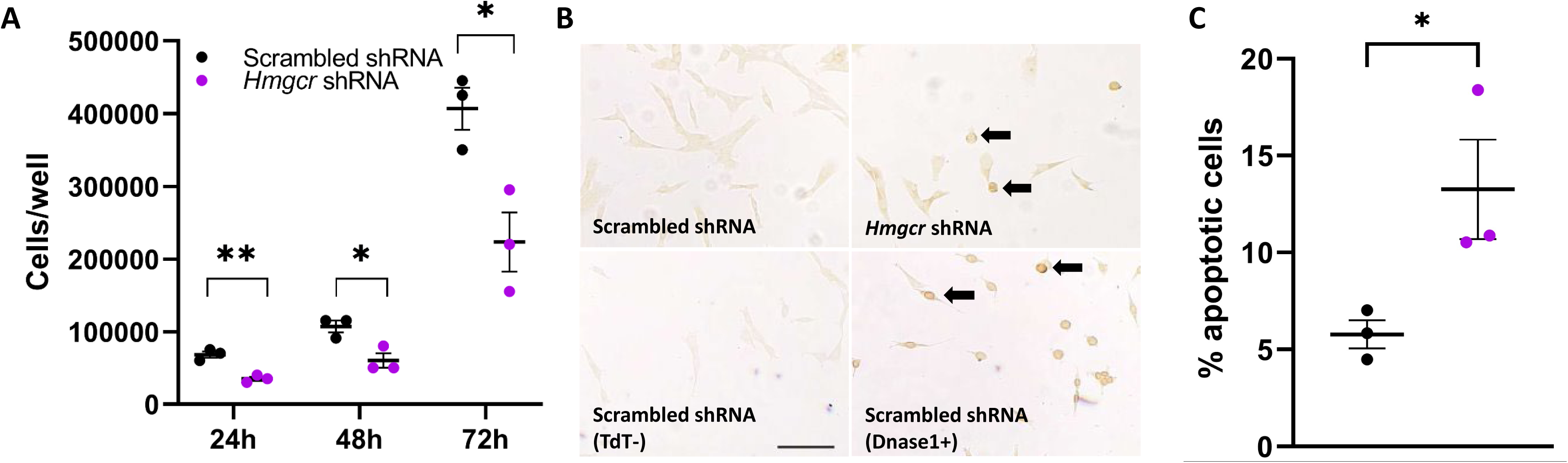
(A) *Hmgcr* shRNA and scrambled shRNA control cells were seeded and counted daily for 3 days. *Hmgcr* knockdown cells showed decreased proliferation compared to scrambled controls. Unpaired t-tests showed p values as follows: **, p= 0.0032 (day 1); *, p=0.0214 (day 2); and *, p=0.0211 (day 3). N = 3 experiments. (B) Dead End Calorimetric TUNEL assay was used to detect apoptotic cells, identified as dark stained nuclei indicating nuclear fragmentation and condensation. Dnase 1 treatment served as a positive control, and absence of the TdT enzyme mix served as a negative control. *Hmgcr* knockdown is associated with an increased number of apoptotic cells compared to scrambled shRNA control (black arrows). Representative microscopic images of n=3 experiments are shown. Scale bar, 100μm. (C) Quantification of the TUNEL assay was conducted via manual counting of dark brown clumped cells. An unpaired t test showed *, p=0.0488.

### Myoblast differentiation in Hmgcr deficiency

Upon culturing in differentiation media for 7 days, *Hmgcr* knockdown myoblasts showed reduced differentiation on days 3 and 7 (Figure 4a) with downregulated *MyoG* gene expression levels compared to scrambled shRNA controls (Figure 4b). Desmin staining (Figure 4c) and myotube fusion indices (Figure 4d) indicated impaired myotube formation in the setting of *Hmgcr* knockdown compared to scrambled controls.

**Figure 4.**
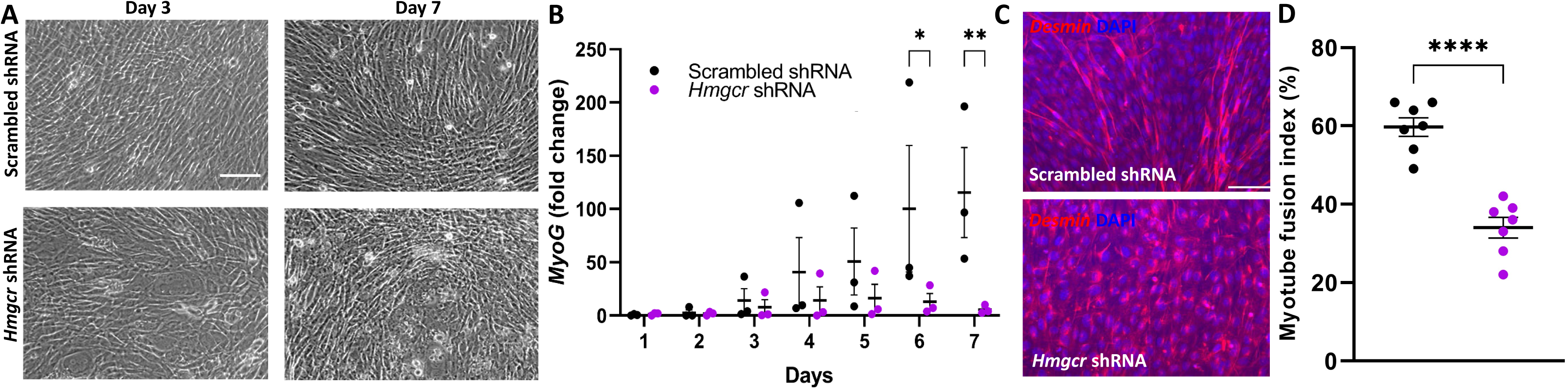
(A) *Hmgcr* knockdown and scrambled shRNA control myoblasts were seeded and switched to differentiation medium at ∼80% confluence. *Hmgcr* deficient cells showed decreased differentiation potential at days 3 and 7, shown in representative images. Scale bar, 100µm. (B) The graph shows significant downregulation of *MyoG* gene expression levels normalized to *Gapdh* in *Hmgcr* knockdown cells compared to scrambled control cells in n=3 experiments. *ANOVA* analysis showed *, p=0.0416; **, p=0.0078. (C) *Hmgcr* shRNA and scrambled shRNA control myoblasts were differentiated and stained for desmin. *Hmgcr* knockdown cells showed impaired myotube formation compared to scrambled shRNA control cells. Scale bar, 100µm. (D) The myotube fusion index from n=3 independent experiments was significantly lower for *Hmgcr* knockdown cells compared to scrambled shRNA controls. An unpaired t-test showed ****, p <0.0001.

### Identification of differentially expressed genes in Hmgcr knockdown myoblasts

We performed nanopore long read transcriptome sequencing on cDNA reverse transcribed from RNA representing *Hmgcr* knockdown vs scrambled shRNA control myoblasts. Deseq2 analysis yielded a total of 299 significant differentially regulated genes in *Hmgcr* knockdown cells compared to scrambled shRNA control cells (Figure 5a and Table S1). There were 190 upregulated genes, including the following genes expressed in skeletal muscle: *Notch3* (Neurogenic Locus notch homolog Protein 3)*, Des* (Desmin)*, Myl9* (Myosin Light peptide 9), and *Scn5a* (Sodium Channel protein type 2 subunit alpha), along with the mitochondrial genes *mt-cytb* (Cytochrome b)*, Ogdhl* (Oxoglutarate dehydrogenase L), and *Nptx1* (Neuronal Pentraxin 1) (Table 1). The cardiac related genes *Sema3c* (Semaphorin-3C) and *Angpt1* (Angiopoietin 1) were also upregulated, as well as *Ckb* (brain specific creatine kinase)^19^. Among the significant differentially expressed genes, 109 genes were downregulated, including the following genes expressed in skeletal muscle: *Myo1b* (Myosin1B)*, Mbnl1* (Muscleblind-like Splicing regulator1)*, Myl6* (Myosin Light polypeptide6), and *Apln* (Apelin) (Table 2). Other downregulated genes include mitochondrial genes whose protein products are involved in enzymatic function and energy metabolism such as *Gtbp10* (GTP Binding Protein 10)*, Gpt2* (Glutamic-Pyruvic Transaminase 2)*, mt-Co1* (Cytochrome C Oxidase 1), and *Acadm* (Acyl-Coenzyme A Dehydrogenase). The proliferation marker *Mki67* was also downregulated in *Hmgcr* knockdown cells compared to scrambled shRNA control cells, which is consistent with our finding of diminished myoblast proliferation.

**Figure 5.**
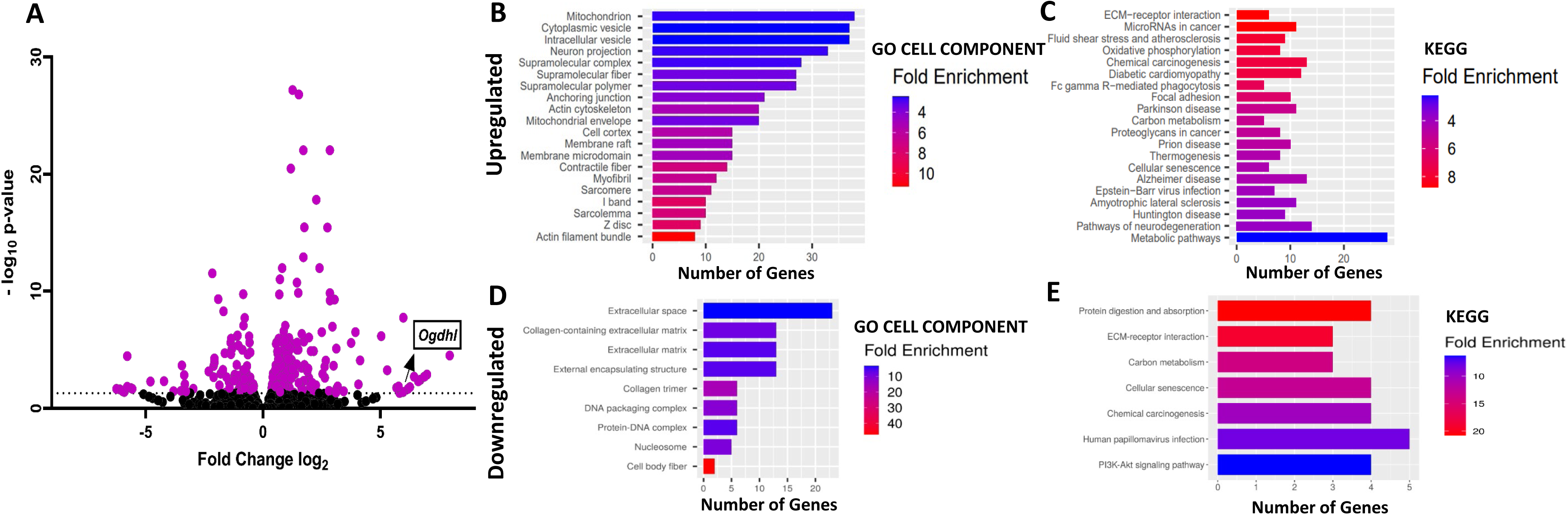
(A) Total RNA was isolated from *Hmgcr* shRNA knockdown and scrambled shRNA cells. Reverse transcription was conducted to generate cDNA libraries for 2 biological replicates of nanopore sequencing for each condition. A total of 299 significant differentially expressed genes were identified using Deseq2. These genes are represented in a volcano plot with the *Ogdhl* gene highlighted. The dotted line indicates p-value=0.05. (A). The outliers *Nefl*, *Il33*, *Prl2c3*, *Myl9*, *Stmn2*, *Peg10* and *Crabp2* were excluded from the volcano plot to facilitate visualization of the core pattern. Among the 299 genes differentially expressed between *Hmgcr* shRNA knockdown and scrambled shRNA control cells, 109 genes were downregulated and 190 were upregulated genes. These genes were analyzed using ShinyGO 0.77. Gene ontology categories are shown based on the (B) cellular components and (C) KEGG pathways for upregulated genes, and the (D) cellular components and (E) KEGG pathways for downregulated genes.

**Table 1.** Selected upregulated mitochondrial and muscle genes.

|  | Upregulated Genes | P value (Adj.) | Fold change |
| --- | --- | --- | --- |
| **Mitochondria** | Ckb | 3.68E-03 | 104.16 |
|  | Ogdhl | 2.21E-02 | 72.07 |
|  | As3mt | 8.18E-05 | 17.58 |
|  | Nptx1 | 5.15E-10 | 8.26 |
|  | Sqor | 2.86E-02 | 4.67 |
|  | Anxa6 | 5.79E-03 | 1.55 |
|  | Prdx5 | 2.17E-02 | 1.42 |
|  | Vat1 | 4.54E-02 | 1.39 |
|  | Cox6b1 | 4.38E-02 | 1.332 |
|  | mt-Cytb | 2.20E-03 | 1.33 |
|  | Vdac2 | 1.33E-03 | 1.48 |
| **Muscle** | Myl9 | 2.14E-34 | 37.91 |
|  | Igfbp5 | 6.85E-07 | 32.80 |
|  | Scn5a | 2.62E-06 | 13.48 |
|  | Npnt | 6.11E-10 | 7.22 |
|  | Chrna1 | 1.05E-12 | 5.30 |
|  | Notch3 | 1.07E-02 | 4.76 |
|  | Plau | 1.30E-05 | 4.46 |
|  | Cdh15 | 1.30E-05 | 4.11 |
|  | Des | 8.85E-08 | 1.92 |

**Table 2.** Selected downregulated mitochondrial and muscle genes.

|  | Downregulated Genes | P value (Adj.) | Fold change |
| --- | --- | --- | --- |
| **Mitochondria** | Gtpbp10 | 3.47E-02 | 0.015 |
|  | G0s2 | 2.43E-04 | 0.232 |
|  | Gpt2 | 4.54E-02 | 0.332 |
|  | mt-Co1 | 1.79E-02 | 0.372 |
|  | Acadm | 1.94E-04 | 0.460 |
|  | Arl2bp | 2.85E-02 | 0.559 |
|  | Ppid | 2.64E-02 | 0.628 |
|  | Mrpl17 | 2.35E-02 | 0.653 |
|  | Slc25a4 | 7.16E-07 | 0.663 |
| **Muscle** | Gldn | 3.44E-02 | 0.015 |
|  | Myo1b | 4.02E-02 | 0.016 |
|  | Cnn1 | 3.47E-02 | 0.071 |
|  | Apln | 1.21E-04 | 0.238 |
|  | H2-K1 | 5.60E-05 | 0.238 |
|  | mt-Co1 | 1.79E-02 | 0.372 |
|  | Ptgfrn | 2.50E-02 | 0.486 |
|  | Kirrel | 4.01E-02 | 0.533 |
|  | Csrp2 | 4.51E-03 | 0.551 |
|  | Mbnl1 | 2.47E-04 | 0.623 |
|  | Mgp | 2.47E-03 | 0.646 |
|  | Myl6 | 1.71E-02 | 0.755 |

### Categories of differentially expressed genes

For the *Hmgcr* knockdown myoblasts compared to scrambled shRNA control myoblasts, gene ontology analysis showed that numerous upregulated genes are related to mitochondrial function (Figure 5b). KEGG pathway analysis showed that genes related to metabolic pathways were most abundantly upregulated (Figure 5c). Genes expressed in the extracellular matrix were represented among the downregulated genes, potentially related to the morphological abnormalities we observed in the *Hmgcr* knockdown myoblasts (Figure 5d). Genes related to cellular senescence were downregulated (Figure 5e), which may in part explain the changes observed in apoptosis patterns (Figure 3b-c).

### Oxygen consumption rate (OCR) analyses

In response to the observed upregulation of numerous mitochondrial genes on the cDNAseq analysis, we found that OCRs were higher in *Hmgcr* shRNA myoblasts compared to scrambled controls (Figure 6a-c). Rotenone treatment confirmed that the increased oxygen consumption was linked to changes in mitochondrial respiration (Figure 6d). A 3-(4,5-dimethylthiazol-2-yl)-2,5-diphenyltetrazolium bromide (MTT) assay showed increased mitochondrial succinate dehydrogenase (complex II) activity in *Hmgcr* shRNA knockdown myoblasts compared to scrambled shRNA controls (Figure 6e).

**Figure 6.**
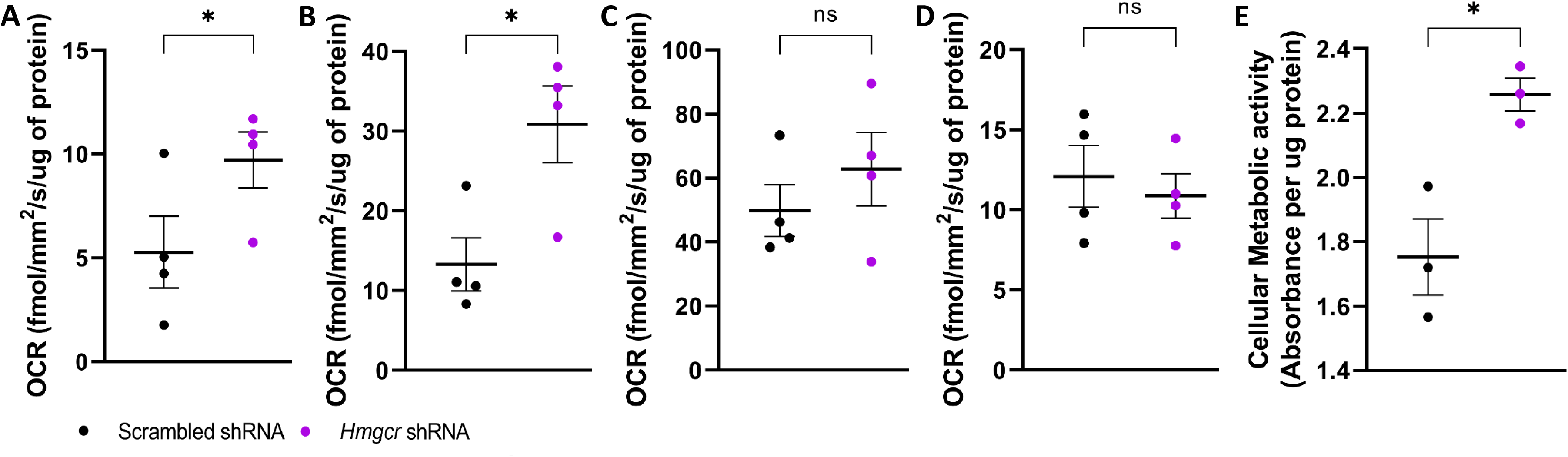
The oxygen consumption rate (OCR) of *Hmgcr* shRNA versus scrambled shRNA C2C12 myoblasts was determined using the RESIPHER system at (A) 24 hours, (B), 48 hours, and (C) 72 hours. The OCR of the *Hmgcr* knockdown cells was increased compared to scrambled controls at 24 and 48 hours. (D) Rotenone treatment at 72 hours confirms that the increased oxygen consumption occurred via mitochondrial respiration. (E) A A 3-(4,5-dimethylthiazol-2-yl)-2,5-diphenyltetrazolium bromide (MTT) assay at 72 hours shows that the total cellular metabolic activity is higher in *Hmgcr* knockdown cells compared to scrambled controls. Paired t-tests were performed. *, p <0.05. Error bars show standard error of the mean (SEM).

### Effects of RNAi-mediated knockdown of hmgcr (CG10367) in Drosophila

Statin administration in flies leads to motor difficulties^20^, indicating that fruit flies are effective models of SAM and potentially other HMGCR-associated muscle diseases. Of note, the *HMGCR* p.Arg443 and p.Ala541 residues impacted by two of the pathogenic variants in the current study are conserved in Drosophila (Figure S2). RNAi techniques were used to knockdown *hmgcr* in Drosophila (Figure 7a). Ubiquitous downregulation of *hmgcr* was lethal during development, whereas *hmgcr* overexpression was well-tolerated (Figure 7b). The lethality phenotype was also observed when downregulating *hmgcr* at the differentiation stage of myogenesis, using the *Mef2-Gal4* driver (Figure 7c).

**Figure 7.**
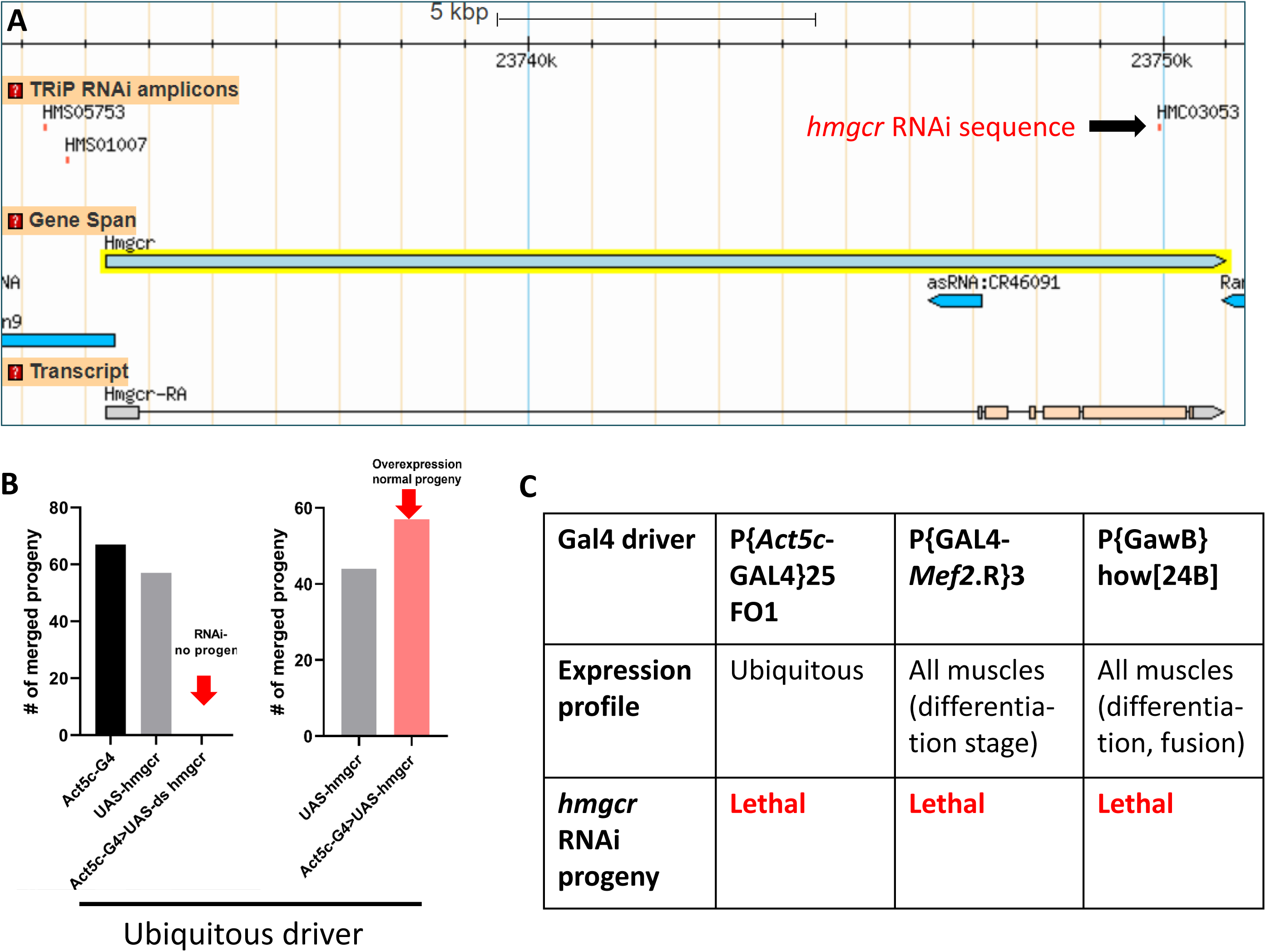
RNAi mediated-knockdown of the homologous fly gene *hmgcr* enables identification of the tissues that are sensitive to loss of function. (A) The *hmgcr* RNAi sequence is aligned on the *hmgcr* locus (arrow, FlyBase, GBrowse). Corresponding mutant fly lines were obtained. (B) On the left, ubiquitous downregulation of *hmgcr* (*Act5C-G4>UAS-ds hmgcr* Drosophila) leads to lethality while two sets of *hmgcr^+/+^*control (Cy) siblings emerge normally (*Act5c-G4* and *UAS-hmgcr*). Conversely on the right, ubiquitous *hmgcr* overexpression (*Act5C-G4>UAS hmgcr* Drosophila) is well tolerated. RNAi and overexpression are accomplished via the Gal4/UAS binary system. Ds, double stranded; Act5C, cytosolic Actin5C. (C) The lethality phenotype is recapitulated when *hmgcr* knockdown is under the control of either the muscle Gal4 driver *Mef2* (number of emerged progeny: control flies n=80, experimental RNAi flies n=0, one replicate), or the muscle Gal4 driver *how* (# of emerged progeny: control flies n=101, experimental RNAi flies n=0, three replicates).

### Effects of HMGCR variant expression on myoblast proliferation and differentiation

Reference sequence human *HMGCR* cDNA and human *HMGCR* cDNA expressing the variants p.Arg443Trp, p.Ser508del, and p.Ala541Thr were stably overexpressed in *Hmgcr* knockdown C2C12 myoblasts and scrambled shRNA control myoblasts. Expression of reference sequence human *HMGCR* showed moderate but inconsistent rescue effects on myoblast proliferation (Figure 8a) and MyoG expression levels (Figure 8b), with more robust and consistent rescue of the myotube fusion index (Figure 8c). Expression of the three variant HMGCR constructs showed attenuated rescue effects (Figure 8a, b, c).

**Figure 8.**
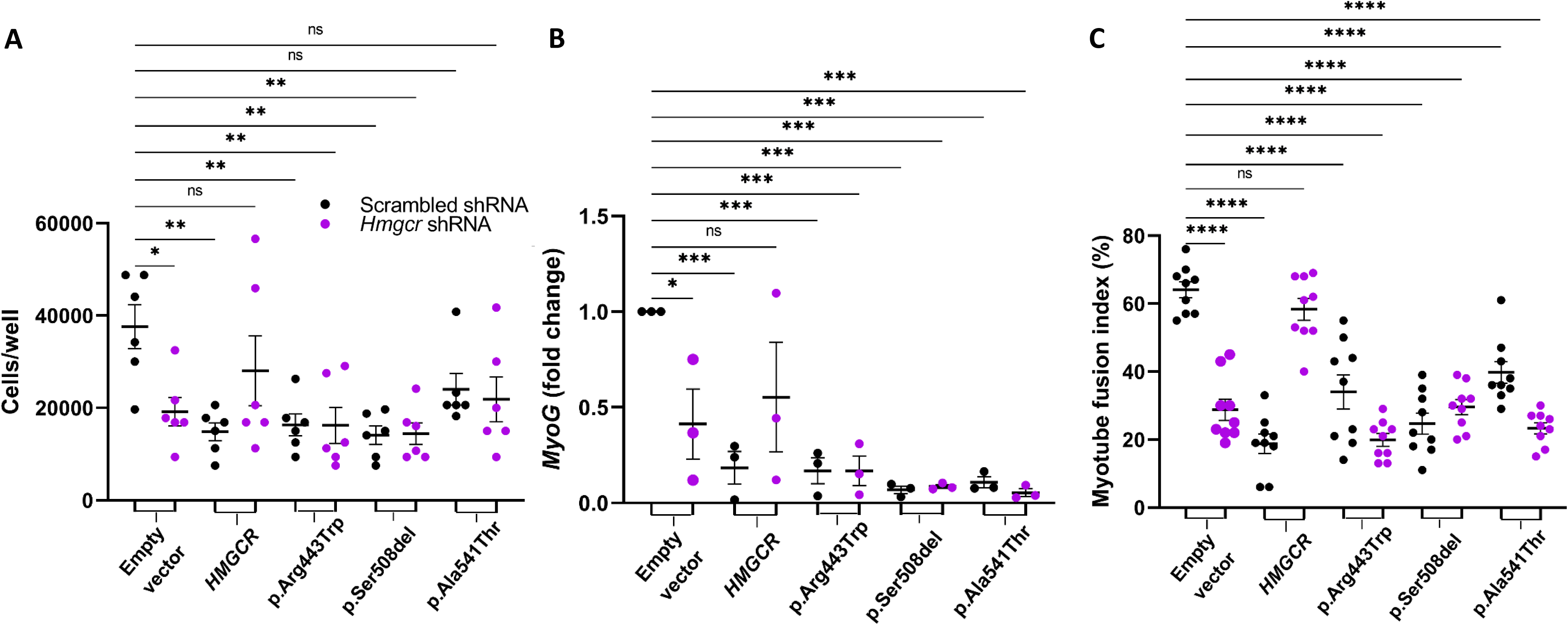
Human reference *HMGCR* and 3 pathogenic variants of *HMGCR* were stably expressed in *Hmgcr* knockdown and scrambled shRNA cells. (A) On day 3 of culture, there were moderate rescue effects with expression of reference *HMGCR* but not with the variant *HMGCR*s on proliferation patterns of *Hmgcr* knockdown cells. N=6 experiments. (B) On day 7 of differentiation, qPCR showed moderate rescue of mRNA *MyoG* expression in *Hmgcr* knockdown cells with expression of reference *HMGCR* but not with expression of the 3 variant *HMGCR*s. N=3 experiments. (C) On day 4 of differentiation, the myotube fusion index in *Hmgcr* knockdown cells showed recovery with expression of reference *HMGCR* but not with the 3 variant *HMGCR*s. *, p < 0.05; **, p < 0.01; ***, p < 0.001; ****, p < 0.0001. *ANOVA* was conducted throughout. Error bars show standard error of the mean (SEM).

## Discussion

Our study shows multi-faceted effects of *Hmgcr* deficiency on myoblast proliferation and differentiation, as well as impaired viability of Drosophila, with impaired rescue effects on myoblasts in the setting of variant *HMGCR* expression compared to reference sequence *HMGCR*. These findings indicate that *Hmgcr* deficiency and pathogenic variants in *HMGCR* are associated with impaired skeletal muscle development, consistent with the childhood onset of several reported cases of HMGCR-related muscular dystrophy^16^.

Statins induce apoptosis in multiple muscle cell types, including skeletal myoblasts^16^, myotubes^21^, and mature myofibers^22^. Muscle regeneration is impaired by statin therapy^23^. Multiple statins, including atorvastatin, simvastatin, and cerivastatin, cause major skeletal muscle-related side effects including rhabdomyolysis, which has been attributed to their lipophilic nature^24^. This hints at the possibility that disruption of cell membrane integrity may contribute to skeletal muscle toxicity in HMGCR deficiency. These effects appear to be specific to skeletal muscle, though cardiac muscle might also be susceptible to detrimental effects of HMGCR deficiency^25,26^.

The Deseq2 analysis on myoblasts highlighted significant differentially expressed genes in sarcomeric binding and energy metabolism. The numerous sarcomeric myosin and actin filament binding genes that were dysregulated in the setting of *Hmgcr* knockdown indicate the importance of HMGCR in maintaining sarcomeric structure. These findings may account for the morphological alterations in the subset of our *Hmgcr* deficient cells showing impaired structure, disorganized nuclei, and loss of membrane integrity. Certain sarcomeric myosin genes were upregulated in the *Hmgcr* knockdown condition indicating potential differences in myofiber contractility. Consistent with our observations, shortened sarcomeres have been noted in a zebrafish model of statin myopathy^27^. In the setting of Hmgcr knockdown, *Des* expression was upregulated at the myoblast stage whereas desmin staining showed downregulation in myotubes, differing from the pattern typically seen during these stages of myogenesis^28^ and indicating that aberrant *Des* expression may also contribute to our structural findings.

Mitochondrial dysfunction has been reported for all three muscle diseases associated with HMGCR. Skeletal muscle histology of skeletal muscle-specific *Hmgcr* knockout mice showed mitochondrial impairment^18^. Knockdown of *hmgcr* in mature myofibers in Drosophila using the *Mhc-Gal4* driver results in viable flies with decreased locomotor activity and fragmented mitochondria^20^, indicating that hmgcr deficiency is consequential both during development and in mature flies, with a more severe phenotype when knocked down earlier in life. SAM is associated with mitochondrial oxidative stress^29^ and mitochondrial dysfunction has been noted on muscle biopsy of a human with anti-HMGCR immune-mediated myopathy^30^. Our cDNAseq data, along with the OCR and MTT findings, demonstrate that the cellular phenotype of *Hmgcr* knockdown is accompanied by elevated mitochondrial respiration. This may be a compensatory response to the *Hmgcr* deficiency. The protein product of the upregulated mitochondrial gene *Ogdhl* mediates the decarboxylation of alpha-ketoglutarate to succinyl-coenzyme (Co)A in the tricarboxylic acid (TCA) cycle^31,32^. The reaction also yields NADH that stimulates NADH dehydrogenase or complex I of the electron transport chain (ETC). Simultaneously, in the TCA cycle, succinyl-CoA is catalyzed by succinyl-CoA synthase to generate succinate which is used by succinate dehydrogenase or complex II of the ETC. Hence energy metabolism pathways may offer future therapeutic strategies for HMGCR-associated muscle diseases and other types of muscle diseases involving the mevalonate pathway.

Other inherited muscle diseases involving the mevalonate pathway include those associated with variants in *HMGCS1* related to rare form of rigid spine syndrome^33^ and those associated with variants in *GGPS1*, which is associated with a rare form of muscular dystrophy represented with hearing loss and ovarian insufficiency^34,35^. HMG-CoA synthase 1 (encoded by *HMGCS1*) catalyzes a reaction immediately upstream of HMGCR, whereas geranylgeranyl diphosphate synthase 1 (encoded by *GGPS1*) catalyzes a reaction in a different downstream pathway than cholesterol synthesis^5^, suggesting that the myopathic effects of HMGCR deficiency may be due in part to protein geranyl geranylation (Figure 1). The classic fibrate medications enhance peroxisome proliferation activator receptor alpha (PPARα) levels, stimulating lipid metabolism. Fibrates are used in combination with statins and are also known to cause myopathy^36^.

It is not clear whether there is a direct connection between cholesterol metabolism and the muscle developmental abnormalities seen in our study. The deleterious effect of inactivation or decreased activity of HMGCR in skeletal muscle raises the possibility that decreased cholesterol levels may contribute to muscle disease. We did not examine this question directly in the current study, but it would merit examination in the future. Abnormal lipid profiles have been observed in humans with dystrophinopathy^38^ and dysferlinopathy^39^. Statin therapy does not appear to be a viable primary or adjunctive therapy for muscular dystrophy, as confirmed in multiple mouse studies^39–42^. However, the non-statin cholesterol-lowering agent ezetimibe showed promise in mouse models of dystrophinopathy and dysferlinopathy^43^. Increased lipids are also not a solution, as they exacerbate the muscular dystrophy phenotype^44,45^. These studies indicate that direct manipulation of lipid levels does not clearly show promise, at this time, as a therapeutic strategy for muscular dystrophy.

Our myoblast culture studies, including transcriptome analysis, in the context of the background literature, suggest that effects on sarcomeric integrity, sarcolemmal integrity, and mitochondrial function drive the myotoxic effects of HMGCR depletion more than decreased cholesterol synthesis. Therapeutic strategies for HMGCR-related muscle diseases would be more likely to be efficacious if they have beneficial impacts on sarcomere/sarcolemmal structure and mitochondrial function.

## Materials and Methods

### Cell culture

C2C12 myoblast cultures were established and maintained in growth medium containing DMEM (*Corning*) +10% fetal bovine serum (FBS) (*Gen Clone*) + 1% penicillin/streptomycin (*Gibco*). To induce differentiation, the cells were then cultured in low serum medium supplemented with DMEM + 2% horse serum (*Gibco*). The differentiation medium was replenished every 48 hours.

### shRNA mediated gene knockdown

C2C12 myoblasts were transfected with either a cocktail of three different shRNAs targeting mouse-*Hmgcr* or scrambled shRNA control (*GeneCopoeia*) using Lipofectamine 3000 (*Invitrogen*). The 3 shRNAs target the following sequences on mouse *Hmgcr* – NM_008255.2:

*Hmgcr* (1) – GGTTGACGTAAACATTAACAA

*Hmgcr* (2) – GGAAACTCTAATGGAAACTCA

*Hmgcr* (3) – GGACATTGAGCAAGTGATTAC

Stable cells expressing GFP reporter were selected using puromycin 3µg/mL (*Sigma – P8833*). GFP expression was detected using Leica DM IL LED microscope (Leica). The GFP-positive clones were expanded and used for further experimentation.

### RNA isolation and qPCR

Total RNA was isolated from shRNA treated cells using an RNA isolation kit (*Zymo research*). Reverse transcription of mRNA was performed using a high-capacity RNA to cDNA kit (*Applied Biosystems*). qPCR-based gene expression analysis was conducted using the Taqman Fast Advanced Master Mix in the QuantStudio 3 Real-Time PCR System *(Thermo Fisher Scientific)* using the following Taqman probes (*Thermo Fisher Scientific*): mouse *Hmgcr* (Mm01282499_m1); mouse *Myod* (Mm00440387_m1); mouse *Myog* (Mm00446194_m1); mouse *Gapdh* (Mm99999915_g1); and human *HMGCR* (Hs00168352_m1).

### Quantification of cell proliferation

shRNA treated cells were seeded and suspended in DMEM medium with 0.4% trypan blue solution at a 1:1 dilution. The cells were then trypsinized and resuspended in growth medium. Viable cells were then manually counted using a hemocytometer on days 1, 2 and 3 using the EVOS XL core microscope (*Life Technologies*).

### Terminal deoxynucleotidyl transferase dUTP nick end labeling (TUNEL) assay

A DeadEnd Colorimetric TUNEL assay kit (*Promega - G7360*) was used to detect apoptosis in *Hmgcr* knockdown cells. Scrambled and *Hmgcr* shRNA knockdown C2C12 myoblasts were seeded at a density of 10,000 cells per well on coverslips. The following day, the TUNEL assay was performed following the manufacturer’s protocol. The slides were imaged using a DM750 microscope (*Leica*). Quantification was performed by manual counting of brown stained (apoptotic) cells.

### Myotube fusion index

C2C12 myoblasts were grown in normal growth medium containing 10% FBS and 1% penicillin-streptomycin on coverslips to 80% confluence. Then the cells were switched to differentiation medium with 2% horse serum. On day 4, the cells were fixed in 4% paraformaldehyde (PFA) for 15 minutes and blocked in serum medium for 1 hour. The cells were stained with Rb-Desmin (*Abcam-ab8592*) or MHC (*DSHB-MF20*) primary antibody for 1 hour. Cells were then stained with anti-rabbit or mouse Alexa Fluor-568 secondary antibody for 1 hour. Nuclei were stained using DAPI and the coverslips were mounted using Fluoromount Aqueous Mounting Medium (*Sigma-Aldrich*). The slides were imaged using a DM6000B or DM5500B epifluorescent microscope (*Leica*). The myotube fusion index was determined by counting the number of nuclei within MHC-positive myotubes divided by the total number of nuclei in the field of view using ImageJ (*National Institutes of Health*).

### Drosophila studies

All strains were raised at 25°C in a 12h light / 12h dark cycle on standard Drosophila media. Genetic crosses were set up at 29°C to generate both the experimental progeny and control siblings under study (the higher temperature increases the efficiency of the yeast Gal4/UAS bipartite expression system), as previously described^46^.

### Human subjects

Clinical and genetic data on study participants were collected and analyzed in accordance with protocols approved by the Institutional Review Board (IRB) or Ethics Committee at the Hospital Sant Joan de Déu, Boston Children’s Hospital, the National Institutes of Health, and the University of Minnesota. Written informed consent was obtained for all study participants.

### Site-directed mutagenesis

Site-directed mutagenesis was performed on human *HMGCR* transcript 1 coding sequence in the pCMV-SPORT6 backbone to generate the variants of interest using the Q5 Site-Directed Mutagenesis Kit (*New England Biolabs*). pCMV-SPORT6-hHMGCR1 was a gift from Anne Galy (Addgene plasmid # 86085; http://n2t.net/addgene:86085; RRID:Addgene_86085). Mutagenic primers harboring the desired variants were used in a PCR reaction with wild type human *HMGCR* sequence as the template. PCR was performed using the following primers for the indicated *HMGCR* variants:

R443W-Forward: 5’ CAGGGAACCTtGGCCTAATGA 3’

R443W-Reverse: 5’ GGAAGTTCAATTTCAGGTTC 3’

S508del-Forward: 5’ CTCCAGTACCTACCTTAC 3’

S508del-Reverse: 5’ AGAAGGTTCTGAAAGCTTC 3’

A541T-Forward: 5’ TGTTGGAGTGaCAGGACCCCT 3’

A541T-Reverse: 5’ GGGATGGGCATATATCCAATAAC3’

The PCR products, which were the linear mutated human *HMGCR* sequences in the same plasmid backbone, were then treated with the provided kinase, ligase, and DpnI enzyme mix to circularize the plasmid, and then transformed into 5-alpha competent *E. coli* (*New England Biolabs*). After transformation, colonies were picked, plasmid DNA was extracted and Sanger sequenced to verify that the desired variant was present. The *HMGCR* sequences were then cloned into pcDNA3.1+ for downstream experiments.

### Stable overexpression of HMGCR variants in C2C12 myoblasts

*Hmgcr* shRNA knockdown and scrambled shRNA C2C12 myoblasts were transfected using Lipofectamine 3000 (*Invitrogen*) with pcDNA3.1+ encoding control *HMGCR* or the 3 *HMGCR* variants of interest: p.Arg443Trp, p.Ser508del, and p.Ala541Thr, as well as empty pcDNA3.1+ controls. Selection of cells expressing the variants was performed using Geneticin 2mg/ml (*Roche*). Stable overexpression of the variants was confirmed via qPCR using human *HMGCR* Taqman probes.

### Nanopore cDNA sequencing and differential gene expression analysis

RNA was extracted from two biological replicates of scrambled shRNA C2C12 cells and two biological replicates of *Hmgcr* knockdown shRNA C2C12 cells using the PureLink RNA Mini Kit (*Invitrogen*). 1μg of the extracted RNA was used as input for the Nanopore cDNA library prep. Nanopore cDNA libraries for the 4 RNA samples were prepared, barcoded, and pooled into a single library. The ligation sequencing V14 – direct cDNA sequencing (SQK-LSK114) protocol was followed until the elution at the cDNA repair and end-prep step. The cDNA was eluted in 23.5μL water. For barcode ligation, 22.5μL end-prepped cDNA, 2.5μL native barcode (NB01-24 from SQK-NBD114.24), and 25 µL Blunt/TA Ligase Master Mix were combined. 5μL of EDTA was used to inactivate the ligation reaction after 20 minutes of incubation. The remainder of the library prep and loading were conducted per protocol for ligation sequencing amplicons – native barcoding kit 24 V14 (SQK-NBD114.24). The barcoded pooled library was loaded on a PromethION flow cell (FLO-PRO114M), and sequencing was performed on a P2 Solo (*Oxford Nanopore Technologies*). The raw data were basecalled using Guppy (v.6.5.7) with the -- barcode_kits "SQK-NBD114-24" --enable_trim_barcodes flags. The basecalled data for each sample was mapped to the mouse genome (GRCm39) using splice-aware minimap2/2.17, and transcripts were counted with htseq. The transcript counts were input into Deseq2 in R 4.3.0, and significantly differentially expressed genes (padj < 0.05) were recorded.

### Oxygen consumption Rate (OCR)

Oxygen consumption measurements were quantified and compared between *Hmgcr* shRNA and scrambled control C2C12 myoblast cultures in real-time using the Resipher system (Lucid Scientific, Atlanta, Georgia) and a 96-well plate (Nunc, Thermo Fisher Scientific) with a compatible 32 sensor lid. Approximately 2,000 C2C12 cells were seeded per well in rows 3, 4, 9, and 10 where the optical oxygen sensors on the sensor lid are located. Two hours after seeding, the sensor lid was placed on top of the 96-well plate, transferred to the incubator (5% CO2 37°C) and attached to the Resipher system. OCR was measured for a total of 72 hours. At the 72-hour mark, rotenone was added to the wells at a concentration of 1 µM to determine whether primarily mitochondrial mediated oxygen consumption was being measured. To control for cell replication over 24, 48, and 72 hours, protein concentrations were collected from replica plates at the end of each designated time point. Protein concentrations were determined using the DC assay kit (BioRad) according to the manufacturer’s instructions.

### MTT assay

*Hmgcr* and scrambled shRNA cells were seeded at density of 2000 cells per well in a 96 well plate in 100ul growth medium. After 72 hours, 25μL of MTT (5mg/ml, Sigma #M2128) was added to each well and incubated for 2 hours. Then, 100μL DMSO was added and the plate was incubated with shaking for 15 minutes. Then the plate was read in a microplate reader at 570nm. The absorbance values were normalized to total protein amounts.

### Statistics

Statistical analysis was performed using GraphPad Prism 9 software. Two-way *ANOVA* was used for *Hmgcr* knockdown *Myog* and *Myod* graphs, followed by the Bonferroni test. Unpaired t-tests were used to analyze the proliferation, myotube fusion index, and TUNEL assays of the *Hmgcr* knockdown condition. One-way *ANOVA* was used for myotube fusion index assay of pathogenic variants, followed by Fisher’s least significant difference test. All other data were analyzed using one-way *ANOVA,* with repeated measures followed by Fisher’s least significant difference test.

## Acknowledgements

Daniel Natera-de Benito was supported by the Miguel Servet programme from Instituto de Salud Carlos III, Spain (CP22/00141). Carsten Bönnemann was supported by intramural funds from the National Institute of Neurological Disorders and Stroke (NINDS), NIH.

---

Table S1. List of 299 significant differentially expressed genes on transcriptome sequencing (cDNAseq), generated by DESeq2, with fold changes.

**Figure S1.**
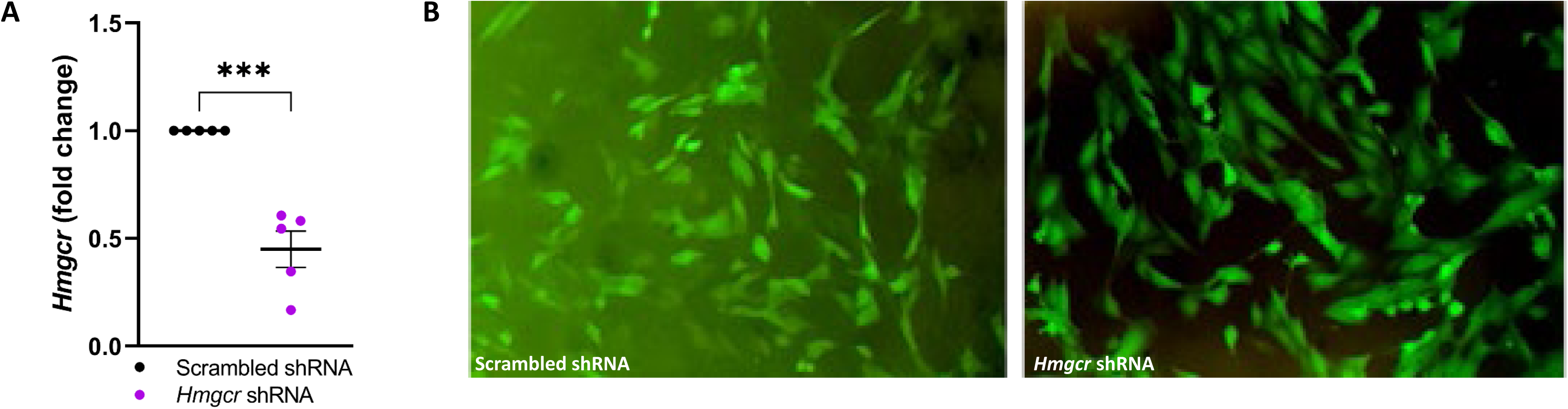
shRNA mediated knockdown of *Hmgcr* in C2C12 myoblasts. (A) qPCR was performed in triplicate for each sample with n=5 experiments. Transcript levels were normalized against *Gapdh* with fold change calculated as *Hmgcr* knockdown versus scrambled control. Unpaired t-test results are shown. ***, p < 0.0002. (B) shRNA mediated knockdown of *Hmgcr* in C2C12 myoblasts. Successful shRNA transfection was confirmed by detecting GFP positive cells.

**Figure S2.**
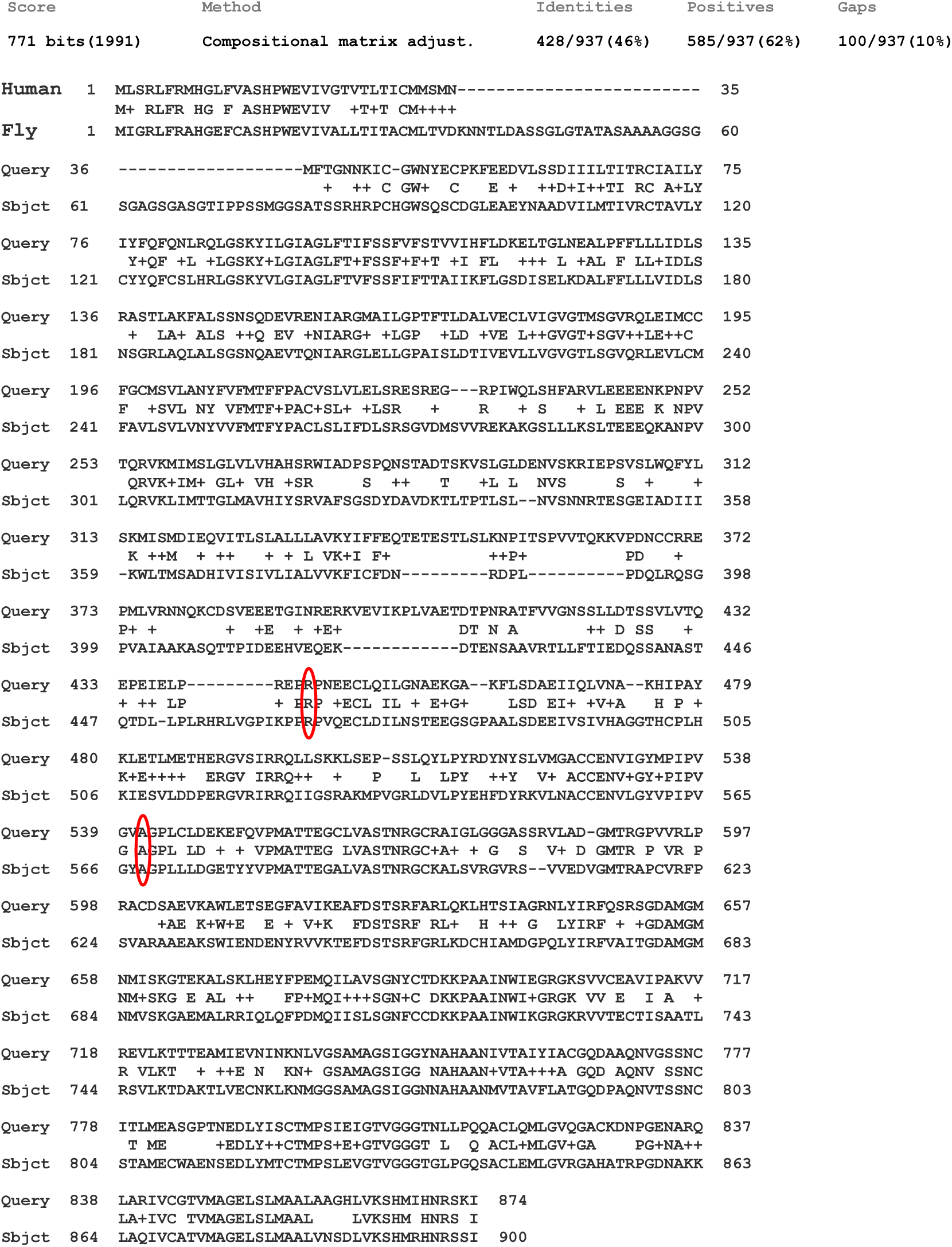
NCBI blastp alignment between Homo sapiens 3-hydroxy-3-methylglutaryl-coenzyme A reductase (*HMGCR*) UniProtKB/Swiss-Prot: P04035 and Drosophila melanogaster *hmgcr* NP_732900.1. The p.Arg443 and p.Ala541 residues targeted by human pathogenic variants are conserved in the fly (red ovals).

